# Stability of the vaginal microbiota during pregnancy and its importance for early infant colonization

**DOI:** 10.1101/2020.04.16.044255

**Authors:** Martin S. Mortensen, Morten A. Rasmussen, Jakob Stokholm, Asker D. Brejnrod, Christina Balle, Jonathan Thorsen, Karen A. Krogfelt, Hans Bisgaard, Søren J. Sørensen

## Abstract

Early life microbiota has been linked to the development of chronic inflammatory diseases. It has been hypothesized that maternal vaginal microbiota is an important initial seeding source and therefore can have lifelong effects on disease risk. To understand maternal vaginal microbiota’s role in seeding the child’s microbiota and the extent of delivery mode-dependent transmission, we studied 700 mother-child dyads from the COPSAC2010 cohort.

The maternal vaginal microbiota was evaluated in the third trimester and compared with the children’s fecal and airway microbiota.

The vaginal samples displayed known stable community state types and only 1:6 changed over time. Only one OTU was significantly transferred to children’s fecal compartment, but an inflated number had positive transfer odds. A few taxonomic families showed early transfer enrichment to vaginally-born children, indicating vertical transfer, while half of the observed transfer effects were delivery mode independent enrichment with attenuating strength over time, indicating a common reservoir.

## Background

Recent studies have suggested, that transfer of bacteria from mother to infant during vaginal birth [1,2] is fundamental for the formation of the early infant microbiota and later disease risk; I) Delivery mode affects the development of the microbiota in early life and differences between the microbiotas of infants delivered vaginally and by cesarean section (CS) can be found up to one year of age [3–5]. II) Such bacterial colonization in early life may influence the risk of several chronic inflammatory disorders [6–10], and III) a clear correlation has been observed between CS and increased risk of such diseases [11,12].

The vaginal microbiota of pregnant women, in comparison with non-pregnant women, has decreased diversity and increased stability [13–17], potentially lowering the risk of bacterial perturbations implicated in adverse pregnancy outcomes, including preterm delivery and low birth weight [18–21], as well as bacterial vaginosis [22]. Several studies of the vaginal microbiota have identified five vaginal community state types (CST); four that are dominated by one *Lactobacillus* species and have low alpha diversity, as well as one containing facultative and strictly anaerobic bacteria, with higher alpha diversity, either dominated by *Gardnerella* spp. or entirely lacking a specific dominating genus [23–31].

While the very first microbial exposure is dictated by delivery mode, mothers’ vaginal microbiota during vaginal birth or skin microbiota after birth by CS [14], the hypothesis of vaginal seeding has been questioned by Wampach et al. which did not observe a difference between the microbiota of vaginal and CS delivered neonates before they were 5 days old [32]. Infants’ airway or fecal microbiota is naturally very different from the vaginal microbiota [2,9,33], and the early development is mainly dependent on various environmental exposures, antibiotic treatments, and genetics [1,3,34–36].

Although the maternal microbiota has been suggested as the most important microbial source for the colonization of the newborn child, previous studies evaluating transfer from mother to child, have not demonstrated strong evidence of transfer, especially not from mother’s vaginal compartment at birth [1]. As a matter of fact Ferretti et al. (2018) shows that bacterial strains from mothers’ stool appear more frequently in the infants’ gut microbiota at a later age [37]. >In this study, we investigated the stability and composition of the vaginal microbiota during the last trimester of pregnancy and its importance for the development of the airway and fecal microbiota from early life up to age three months and one year, respectively. We used samples from the 695 mother-child dyads in the Copenhagen Prospective Studies on Asthma in Childhood 2010 (COPSAC_2010_) cohort from week 24 of pregnancy until one year of age. The bacteria were identified by 16S rRNA gene amplicon sequencing.

## Results

### Vaginal microbiota

The vaginal microbiota at gestational week 24 (n=730) and 36 (n=665) was dominated by bacteria belonging to the three phyla Firmicutes (86.5%), Actinobacteria (10.8%) and Proteobacteria (1.6%). At genus level, this corresponded to *Lactobacillus* (81.9%) and *Gardnerella* (7.8%). The most abundant operational taxonomic units (OTU) were all lactobacilli, with sequences matching *L. crispatus* (33.4%), *L. iners* (25.6%), *L. gasseri* (8.5%), and *L. jensenii* (4.3%). The samples clustered into five CSTs similar to those shown in previous studies (Supplementary Table 1, Supplementary Table 2, and Supplementary Results: <u>Additional file 1</u>). We analyzed the stability of the vaginal microbiota using two separate methods; one based on CSTs and one based on beta diversity.

Of the 657 women with both week 24 and 36 data, 541 (82.3%) had the same CST at week 24 and week 36, with CST IV-b (19 out of 33 - 57.6 %) and CST V (29 out of 43 – 67.4%) being significantly less stable (χ^2^-test, p-value < 10^−5^) (Supplementary Figure 1, Supplementary Table 3; Additional file 1). The median Jensen-Shannon divergence (JSD) between women’s paired week 24 and week 36 samples (0.056) were significantly lower than the median JSD between mismatched pairs of week 24 and week 36 samples (0.617) (p < 10^−3^, Supplementary Figure 2, Supplementary Table 4; Additional file 1). Furthermore, the distance from week 24 to week 36 strongly depends on week 24 CST as women with CST IV-b observed 3.6-7.8 times higher median divergence compared to women with other CSTs (Supplementary Table 5; Additional file 1).

**Figure 1:**
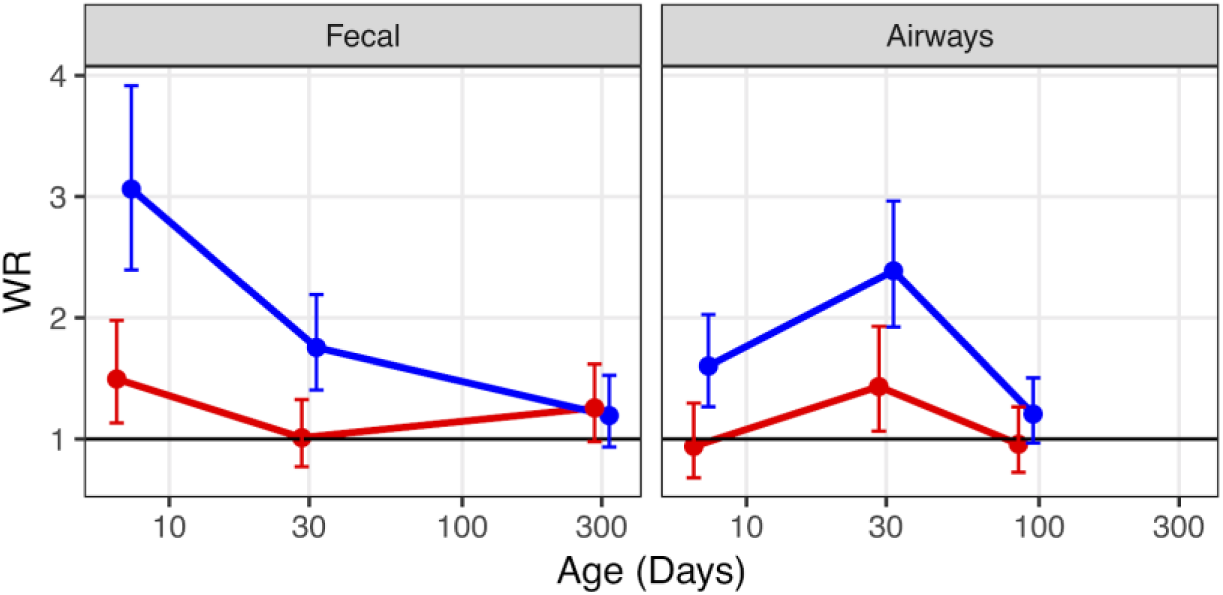
Weighted Transfer Ratios from vaginal week 36 to the fecal- and airway compartment in first year of life stratified on mode of delivery (blue: vaginal birth, red: cesarean section). A ratio above 1 indicates enrichment of microbial transfer. Error bars reflects standard errors

**Figure 2:**
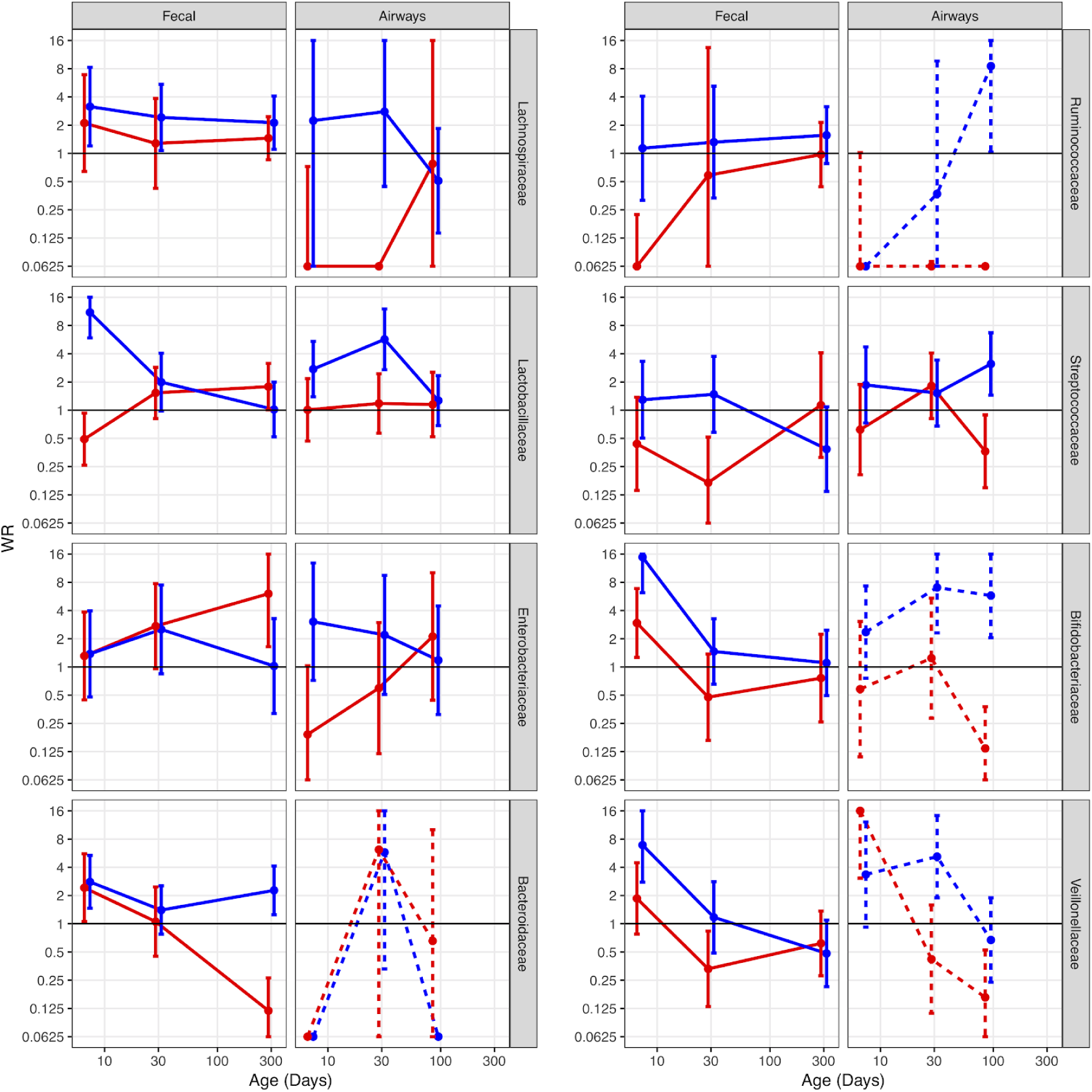
Weighted Transfer Ratios from vaginal week 36 to the fecal- and airway compartment in first year according to mode of delivery (blue: vaginal birth, red: cesarean section) partitioned for the eight most represented taxonomic classes at Family level with upper right (*Lachnospiraceae*) being the most represented family followed by *Lactobacillaceae* and so forth. For compartments with less than 30 representative OTUs, the results are shaded. Error bars reflects standard errors

### Infant microbiota

In the 1,784 fecal samples from the infants (1 week: 552, 1 month: 607 and 1 year: 625) we identified 4,524 OTUs, with high relative abundance of the phyla Bacteroidetes (36%), Proteobacteria (26%), Firmicutes (21%), and Actinobacteria (16%), and in the 1,811 airway samples from the infants (1 week: 544, 1 month: 645 and 3 months: 622) we identified 3,709 OTUs, with high relative abundance of the phyla Firmicutes (61%), Proteobacteria (30%), Actinobacteria (6%), and Bacteroidetes (2%). For a detailed description see Stokholm et al. 2018 and Mortensen et al. 2016 for fecal and airways, respectively. 549 children (78%) were delivered vaginally and 151 children (22%) by CS (of the 700 children enrolled in the study). Delivery mode was associated with the one-week microbiota composition in both airways and especially fecal samples (Bray-Curtis distances PERMANOVA, hypopharynx, p = 0.035 and feces, p < 0.001).

### Transfer of the microbiota

Out of a total of 2,291 identified vaginal OTUs, 352 to 1,182 OTUs were tested for transfer according to compartment and time point. These OTUs covered 33-42%, 39-87% and 34-95% of the vaginal, fecal and airway reads, respectively. Table 1 shows summary statistics on the OTUs tested for each comparison.

**Table 1:** Descriptives on testable OTUs in terms of numbers of OTUs, vaginal, fecal and airway total coverage, number of tests reaching nominal and FDR corrected significance.

| Compartment | Delivery mode | Age | # testable OTUs | Vaginal relative abundance | Child relative abundance | min(p)§ | min(q)* | # p<0.01 | # p<0.05 | # q<0.05 | # q<0.10 |
| --- | --- | --- | --- | --- | --- | --- | --- | --- | --- | --- | --- |
| Fecal | Vaginal | 1 week | 1142 | 41.2 | 61.1 | **4.42E-06** | **5.04E-03** | 20 | 58 | **1** | **3** |
| Fecal | Vaginal | 1 month | 1081 | 40.6 | 63.2 | **2.18E-04** | 2.36E-01 | 15 | 51 | 0 | 0 |
| Fecal | Vaginal | 1 month | 1182 | 42.6 | 87.2 | **1.56E-03** | 9.98E-01 | 2 | 22 | 0 | 0 |
| Fecal | CS | 1 week | 564 | 37.3 | 57.3 | **2.90E-03** | 9.91E-01 | 2 | 11 | 0 | 0 |
| Fecal | CS | 1 month | 569 | 33.2 | 39.0 | **2.56E-02** | 9.91E-01 | 0 | 8 | 0 | 0 |
| Fecal | CS | 1 month | 547 | 36.9 | 58.0 | **4.55E-03** | 9.91E-01 | 2 | 11 | 0 | 0 |
| Airway | Vaginal | 1 week | 764 | 42.4 | 46.7 | **2.83E-03** | 9.98E-01 | 1 | 20 | 0 | 0 |
| Airway | Vaginal | 1 month | 902 | 41.8 | 43.9 | **1.08E-04** | 9.75E-02 | 8 | 29 | 0 | **1** |
| Airway | Vaginal | 3 month | 995 | 42.3 | 73.3 | **9.73E-04** | 9.68E-01 | 5 | 27 | 0 | 0 |
| Airway | CS | 1 week | 352 | 37.9 | 33.9 | **1.98E-02** | 9.90E-01 | 0 | 5 | 0 | 0 |
| Airway | CS | 1 month | 452 | 36.2 | 85.0 | **1.24E-03** | 5.62E-01 | 2 | 10 | 0 | 0 |
| Airway | CS | 3 month | 459 | 36.3 | 95.5 | **1.36E-03** | 6.26E-01 | 1 | 8 | 0 | 0 |

§ uncorrected p-values
\* q-values via FDR correction by Benjamini-Hochberg False Discovery Rate

#### Transfer of specific OTUs

For each testable OTU, we calculated the odds ratio (OR) and p-value for transfer from mothers’ vaginal microbiota to their child’s fecal or airway microbiota at week 1 (Supplementary Figure 3 for vaginally born children and Supplementary Figure 4 for CS-born children: Additional file 1). In general, only a single OTU (Belonging to the family *Lachnospiraceae*) showed false discovery rate (FDR) corrected statistical significant transfer to the fecal compartment. We then examined whether maternal abundances of the OTUs would affect the likelihood of transfer. By correlating the OR for transfer for each OTU with the population wide relative maternal abundance, we revealed that the strongest transfer of OTUs was observed for low abundant vaginal bacteria to the fecal compartment both for vaginally (p < 0.01) and CS-born children (p < 10^−8^), whereas there was no association between population abundance and transfer odds to the hypopharynx (p > 0.2, Supplementary Figure 3 and 4: Additional file 1).

#### Enrichment of OTUs with positive transfer estimates (compared to negative)

In order to pursue an enrichment hypothesis, a weighted transfer ratio between positive- and negative transfer odds was calculated as function of birth mode, compartment and age (Figure 1). For vaginally-born children, a clear enrichment was observed for both compartments at week one (weighted transfer ratio (WTR_VAG_) = 3.6 and WTR_VAG_ = 2.0 for transfer to gut and airways respectively, p < 0.001). For CS-born children a tendency towards enrichment was observed for the fecal compartment at the early time point (WTR_CS_ = 1.7 for transfer, p = 0.06). Transfer to the fecal compartment attenuated with increasing age towards similar results at age one year for both birth modes. Interestingly, transfer to the airways was strongest at age one month for both vaginally and CS-born children (WTR_VAG_ = 2.8 and WTR_CS_ = 1.8, with p < 0.001 and p = 0.04 for vaginally and CS-born children respectively).

A detailed enrichment analysis, grouping OTUs at family level for the eight most common bacterial families (*Lachnospiraceae, Lactobacillaceae, Enterobacteriaceae, Bacteroidaceae, Ruminococcaceae, Streptococcaceae, Bifidobacteriaceae* and *Veillonellaceae*) showed that the overall enrichment result was family dependent (Figure 2). Several patterns were observed, some showed evidence of transfer, where positive and larger WTR are observed for vaginally-born children in comparison with CS-born children at one week, which were either transient (decreased over time) or persistent (maintained over time). Other patterns showed evidence of a common reservoir, where WTR increased in general and became more similar between mode of delivery with increasing age, indicating that bacteria were shared due to common living environments or other external factors, and not transferred from mother to child during birth.

*Lachnospiracea*, the most represented family, shows evidence of persistent transfer to the fecal compartment with positive attenuating associations for vaginally-born children over the first year of life (WTR_vag_ = 3.2 to WTR_vag_ = 2.1 from 1 week to 1 year respectively). *Lactobacillaceae, Bifidobacteriaceae* and *Veillonellaceae* showed strong evidence of transient transfer to the fecal compartment, which rapidly declined at one month of age. A similar trend was observed for transfer to the airways for *Lactobacillaceae* strongest at one month of age (WTR_vag_ = 5.7). Associations between the vaginal and the fecal compartment for *Enterobacteriaceae* and *Bacteroidaceae* likely reflect the existence of a common reservoir, as similar results are obtained for both delivery modes. However, the temporal trends are opposite, such that the associations becomes stronger from one week to one month for *Enterobacteriaceae* (WTR_vag_ = 1.4-2.5 and WTR_CS_ = 1.3-2.7) while the opposite is the case for *Bacteroidaceae* (WTR_vag_ = 2.8-1.4 and WTR_CS_ = 2.4-1.1).

For *Streptococcaceae* and *Ruminococcaceae* there were not observed any clear patterns of associations that could be explained by transient transfer or common reservoir.

#### Transfer of the most dominating vaginal OTU at week 36

For each dyad, the frequency of the most dominating maternal OTU was estimated in the children at one week of age. In the fecal compartment, 30% of CS delivered children and 38% of vaginally delivered children had the most dominating maternal OTU. In the airway compartment, these figures were 16% and 25%. However, a permutation test revealed that this association was only statistically significant for vaginally-born children (p < 0.05 for vaginally and p > 0.2 for CS-born children). These associations attenuate to non-significant for later time points.

## Discussion

The present study has characterized the vaginal microbiota of 730 Danish women during pregnancy and subsequently the microbiota of 695 mother child pairs. In concordance with previous smaller North American and European studies (MacIntyre et al., 2015 (n = 42), Ravel et al., 2013 (n = 396), Romero et al., 2014 (n = 32+22)) [16,29,38] our results confirm in a bigger cohort that the five major CSTs and high stability of the vaginal microbiota in the last trimester of pregnancy. Considering this high stability we utilize the vaginal microbiota at week 36 as a proxy for the infants’ bacterial exposure during vaginal birth.

Just one single OTU demonstrated significant transfer odds from mother to infant after correction for multiple testing, and while we did observe an inflated number of positive correlations (3.6 times more for fecal and 2.0 times more for airway samples) with attenuating strength over the first year of life, the positive correlations were found among most taxa. Interestingly, we found that taxa with lower maternal population abundance were significantly more likely to be transferred. This pattern could indicate that the dominating taxa are highly adapted to their specific microbial niche, and that the species dominating the vaginal microbiota will not necessarily be able to successfully colonize the gut or airways of the child.

Many studies have identified delivery mode as a significant factor for the formation of infants’ early microbiota [1,4,5], and our data confirm this for both airway and fecal samples from one week old infants. We found enrichment of transfer rates for both delivery methods, with the most pronounced correlations to fecal samples being driven by the families *Bifidobacteriaceae* and *Lactobacilliaceae*, for which the larger early enrichment in transfer to vaginally-born children indicated a strong vertical transfer during vaginal birth that attenuated by one month. *Veillonellaceae* had weaker, but similarly developing enrichment over time, while *Enterobacteriaceae* displayed an increase in transfer enrichment to the fecal compartment from one week to one month and continuing for CS-born children suggesting that this merely reflects a shared environment. Similarly, *Bacteroidaceae* reflected an early life shared environment at one week of age, rapidly attenuating at one month of age. That these bacteria are correlated between mother and infant regardless of delivery mode, strongly indicate that their presence in the infant microbiota is not solely due to transfer during birth, but transferred from mother to infant at a later point or caused by the shared living environment of mother and infant. All five bacterial families have been identified as part of the shared microbiota between breast milk, mothers fecal samples and infants fecal samples [34], indicating that microbiota transfer after delivery may be more important than during delivery itself. This is further supported by a study showing that almost 40% of infants’ fecal microbiota in the first 30 days of life originated from either breast milk or areolar skin microbiota [39].

### Conclusion

Our results suggest that mothers’ vaginal microbiota has minor impact on their infants’ airway and fecal microbiota in early life. While infants are exposed to their mothers’ vaginal microbiota, during vaginal birth, the dominating bacteria are not commonly associated with the very different environments in the hypopharynx and intestines. We found that most shared taxa could be attributed to post-delivery transfer, possibly through skin contact, breast milk, or a shared living environment.

## Methods

### Study population

COPSAC_2010_ is an ongoing Danish mother-child cohort study of 700 unselected children and their families followed prospectively from pregnancy week 24 in a protocol previously described [40]. Exclusion criteria were gestational age below week 26; maternal daily intake of more than 600 IU vitamin D during pregnancy; or having any endocrine, heart, or kidney disorders.

### Sample collection

Vaginal samples from the symptom-free women at gestational week 24 and 36 were collected from the posterior fornix of the vagina using flocked swabs (ESWAB regular, SSI Diagnostica, Hillerød, Denmark) [41]. Airway samples were aspirated with a soft suction catheter passed through the nose into the hypopharynx as previously described in detail [6]. Fecal samples were collected in sterile plastic containers and transported (within 24 hours) to Statens Serum Institute (Copenhagen, Denmark). Each sample was mixed on arrival with 10% vol/vol glycerol broth (SSI, Copenhagen, Denmark) and frozen at −80 °C until further processing [9]. 2,670 samples were collected and initially included.

The airway microbiota samples used in this study have been presented previously in Mortensen et al. (2016) [33] and the fecal samples have been presented in Stokholm et al. (2018)[9]. For both sample types, three consecutive samples were included to investigate transfer from mother to infant; from feces at one week, one month and one year, and from airways at one week, one month and three months.

### DNA extraction

Genomic DNA was extracted from the mothers’ and infants’ samples as described in Mortensen et. al (2016) [33], using the PowerMag® Soil DNA Isolation Kit optimized for epMotion® (MO-BIO Laboratories, Inc., Carlsberg, CA, US) using the epMotion® robotic platform model (EpMotion 5075VT, Eppendorf, Hamburg, Germany).

### 16S amplicon sequencing and bioinformatics pipeline

16S rRNA gene amplification was performed as described in Stokholm et. al (2018) [9], using a two-step PCR method, targeting the hypervariable V4 region (Forward primer 515F: 5’-GTGCCAGCMGCCGCGGTAA-3’ [42], reverse primer Uni806R: 5’-GGACTACHVGGGTWTCTAAT-3’ [43]). Amplicon products were purified with Agencourt AMPure XP Beads (Beckman Coulter Genomics, MA, US) and the purified products quantified with Quant-iT™ PicoGreen® quantification system (Life Technologies, CA, US) to allow for pooling, in equimolar concentration, of up to 192 samples per library. The pooled DNA samples were concentrated using the DNA Clean & Concentrator™-5 Kit (Zymo Research, Irvine, CA, US) and quantified again. The libraries were sequenced on the Illumina MiSeq System (Illumina Inc., CA, US) using MiSeq Reagent Kits v2.

The bioinformatics analysis of the demultiplexed Fastq-files were performed using biopieces [44], which wraps around usearch v7.0.1090 (mate-pairing, filtering) [45], UPARSE (OTU clustering at 97% similarity and singleton removal) [46], the UCHIME algorithm (Chimera checking and removal) [47], Mothur v.1.25.0 (taxonomic classification against the Ribosomal Database Project (RDP) trainset9 database (032012) [48]) [49], and FastTree (Creation of a phylogenetic tree) [50]. Sequence contingency tables were exported at OTU level, with taxonomical classification down to genus level if available.

### Identification of lactobacilli

All representative sequences for OTUs assigned to the family *Lactobacilliaceae* were aligned to the NCBI 16S rRNA (Bacteria and Archaea) database using BLAST to assign putative species [51]. The OTUs were assigned to unique species, when possible, but some sequences aligned equally well to two or more species. In these cases, we used published studies to select the most likely species. As an example, *L. crispatus* and *L. acidophilus* could not be distinguished based on the sequenced region and as published studies on the vaginal microbiota concur that *L. crispatus*, in contrast to *L. acidophilus*, constitute an important part of the vaginal microbiota, we refer to *L. crispatus*/*L. acidophilus* solely as *L. crispatus*. Based on this reasoning we will also refer to *L. gasseri/johnsonii*, solely as *L. gasseri* [52–54].

### Bioinformatics analysis

For data treatment and analysis we used the open source statistical program ‘R’ [55], predominantly the R-package “phyloseq” [56]. Samples with less than 2,000 sequences were excluded. 2,359 samples were included containing, on average, over 32,000 sequences per sample, representing 3,934 distinct OTUs. JSD was used to describe the beta diversity in the sample set. As this method is sensitive to bias due to sequencing depth, we performed the calculation of JSD on a randomly subsampled OTU table with an even sequencing depth of 2,000 observations. No other analysis were performed using the subsampled OTU table.

### Clustering analysis

Clustering analysis was performed using hierarchical clustering, based on JSD, and the optimal number of clusters were chosen based on multiple cluster validation techniques giving larger emphasis to OTUs with high inference and effect size. *ORi* and *p.valuei* refers to the OR and its corresponding null hypothesis test respectively for the *i’*th OTU.

## List of abbreviations

COPSAC_2010_: Copenhagen Prospective Study on Asthma in Childhood 2010
CS: Cesarean Section
CST: Community State Type
FDR: False Discovery Rate
JSD: Jensen-Shannon Divergence
OR: Odds Ratio
OTU: Operational Taxonomical Unit
rRNA: ribosomal Ribonucleic Acid
VAG: Vaginal delivery
WTR: Weighted Transfer Ratio

## Declarations

### Ethics approval and consent to participate

This study followed the principles of the Declaration of Helsinki, and was approved by the Ethics Committee for Copenhagen (The Danish National Committee on Health Research Ethics) (H-B-2008-093) and the Danish Data Protection Agency (2008-41-2599). Written informed consent was obtained from both parents for all participants. The study is reported in accordance with the Strengthening the Reporting of Observational Studies in Epidemiology (STROBE) guidelines [61].

### Consent for publication

Not applicable

### Availability of data and material

The dataset analysed during the current study will be available, upon publication, in the Sequence Read Archive (SRA) repository. Before publication reviewers can download all data from https://github.com/mortenarendt/VagTransfer/RawData.zip and decrypt the file with the password VagTransferCOPSAC.

Additional file 1 contains all figures, tables, supplementary figures, supplementary tables, and the R-script for the entire analysis performed in this article, which can be reproduced using the data in COPSACbirthmicobiome.RData (Additional file 2), as well as the two supporting scripts: getTransferStats.R (Additional file 3) and getWinnerStats.R (Additional file 4).

## Authors’ contributions

M.S.M. and M.A.R is the main authors of this paper. M.S.M and C.B performed DNA extraction and sequencing. A.D.B. performed the initial bioinformatics analysis. M.S.M. performed the microbiota analysis. J.S. were responsible for sample collection from mothers and infants. M.A.R and M.S.M performed the statistical data analysis, J.S., J.T. and J.W., helped interpret the data. This project was conceived and designed by H.B., S.J.S., and K.A.K. All the authors have read and understood the manuscript.

## Acknowledgements

We express our deepest gratitude to the children and families of the COPSAC2010 cohort study for all their support and commitment. We acknowledge and appreciate the unique efforts of the COPSAC research team. We thank Susanne Schjørring for critical logistic support, with handling and storing of samples at Statens Serum Institut, Denmark. Furthermore, we thank Karin Pinholt Vestberg, April Cockburn, and Jakob Russel (Section for microbiology, University of Copenhagen) for the help and support with DNA extraction, construction of the 16S rRNA gene amplicon libraries, and sequencing.

## Additional files

Additional file 1: **FullAnalysis.html** is an html file containing the R code for reproducing the entire analysis, including all figures, tables, supplementary figures and supplementary tables.

Additional file 2: **COPSACbirthmicobiome.RData** is an RData file containing all data necessary to reproduce the full analysis for this article.

Additional file 3: **getTransferStats.R** is an Rscript containing additional functions needed to calculate the transfer statistics for all individual OTU models.

Additional file 4: **getWinnerStats.R** is an Rscript containing additional functions needed to complete the calculations of the transfer statistics for the most abundant OTU in each woman’s vaginal microbiome at week 36.

